## Supplemental file 1 for "Proximity proteomics in a marine diatom reveals a putative cell surface-to-chloroplast iron trafficking pathway"

**Table S1:** Predicted endogenous biotinylated *P. tricornutum* proteins are present at similar levels in WT and pTF-APEX2 proteomic samples.

There are at least five predicted biotin-containing proteins in *Phaeodactylum tricornutum*. APEX2/WT ratios near 1 for all of them suggest an unbiased streptavidin pull-down independent of genetic background. Seeing all of them in our dataset also indicates the pull-down itself worked.

| **UniProt ID** | **APEX/WT ratio** | **Ensembl ID** | **Annotation** | **Size [kDa]** |
| --- | --- | --- | --- | --- |
| B7G7S4 | 0.97 | Phatr3_EG01955 | acetyl-CoA carboxylase | 229.5 |
| B7GBG1 | 1.22 | Phatr3_J30519 | pyruvate carboxylase | 136.1 |
| B7GA98 | 0.97 | Phatr3_J49339 | pyruvate carboxylase | 137.9 |
| B7GCL6 | 0.94 | Phatr3_J51245 | propionyl-CoA carboxylase | 72.5 |
| B7GEB5 | 0.89 | Phatr3_J55209 | biotin carboxylase | 249.8 |

**Table S2:** Features of the three proteins co-expressed from a gene cluster on chromosome 20.

| **UniProt ID** | **Ensembl ID** | **Annotation** | **New name*** | **Length**  **[AA]^1^** | **Size**  **[kDa]^1^** | **pI^1^** | **Signal peptide^2^**  **[cut**↓**site]** | **Predicted**  **localization^3^** | **# of transmembrane domains [position]^5^** |
| --- | --- | --- | --- | --- | --- | --- | --- | --- | --- |
| B7G9B3 | Phatr3_J51183 | CREG1-like protein | pTF.CREG1 | 234 | 26.35 | 5.03 | yes [SGA↓YP] | chloroplast^4^ | 0 |
| B7G9B2 | Phatr3_J52498 | cell surface protein | pTF.CatCh1 | 349 | 37.43 | 4.81 | yes [ASA↓EV] | secretory pathway | 1 [296–315] |
| B7G9B0 | Phatr3_J54986 | cell surface protein | pTF.ap1 | 393 | 42.36 | 5.16 | yes [AFS↓VE] | chloroplast^4^ | 1 [327–349] |

*pTF.CREG1: CREG1-like protein; pTF.CatCh1: Chloroplast-AssociaTed metalloCHaperone 1; pTF.ap1: pTF-Associated Protein 1

^1^ProtParam. ^2^SignalP 4.1. ^3^TargetP 1.1, SignalP 4.1., and ASAFind version 1.1.7., ^4^low confidence (as determined by ASAFind), ^5^TMHMM Server v. 2.0.

**Table S3:** Molecular cloning primers.

| **Primer ID** | **Forward [F]**  **or Reverse [R]** | **Sequence**  **[5’ → 3’]** | **Length**  **[nt]** | **Corresponding vector**  **and amplicon details** |
| --- | --- | --- | --- | --- |
| JT01 | F | CGAATCAGGATCTAAAATGAACGCACGTCTGCGACCTGAGCAA | 43 | pJT_NR_pTF-AP2  (backbone) |
| JT02 | R | GTCGCTTCACGTTCGCTC | 18 | pJT_NR_pTF-AP2  (backbone) |
| JT03 | F | GATACGCGAGCGAACGTGAAGCGACTCACGTAGTGAAGTGATGTTG | 46 | pJT_NR_pTF-AP2  (*NR* promoter, 5’ *NR* UTR, *pTF*) |
| JT04 | R | TTCCAGACGTAGAACCACTCCCTTTGATAGGAGTGCTGCCAGTG | 44 | pJT_NR_pTF-AP2  (*NR* promoter, 5’ *NR* UTR, *pTF*) |
| JT05 | F | CGCTCACCGGCTCCAGATTTATCAGCAATAAACCAGCC | 38 | pJT_native_pTF-mCherry  (backbone amplicon 1) |
| JT02 | R | GTCGCTTCACGTTCGCTC | 18 | pJT_native_pTF-mCherry  (backbone amplicon 1) |
| JT07 | F | CGTCTGCGACCTGAGCAAC | 19 | pJT_native_pTF-mCherry  (backbone amplicon 2) |
| JT08 | R | AAATCTGGAGCCGGTGAG | 18 | pJT_native_pTF-mCherry  (backbone amplicon 2) |
| JT09 | F | GATACGCGAGCGAACGTGAAGCGACCCTCGAGCTTGAGTCTCTG | 44 | pJT_native_pTF-mCherry  (*pTF-mCherry* cassette) |
| JT10 | R | CATGTTGTTGCTCAGGTCGCAGACGTCCCTCCTGATACCTACC | 43 | pJT_native_pTF-mCherry  (*pTF-mCherry* cassette) |
| JT11 | F | TAAAGTCTGGAAACGCGGAAG | 21 | pJT_pTF-mCherry_MS hit-EYFP  (backbone amplicon 1) |
| JT12 | R | TTTAAAGAGATCGCAATCTGAATCTTGGTTTCATTTGTAATACGCTTTAC | 50 | pJT_pTF-mCherry_MS hit-EYFP  (backbone amplicon 1) |
| JT13 | F | GATTGCGATCTCTTTAAAGGGTGGTCC | 27 | pJT_pTF-mCherry_MS hit-EYFP  (backbone amplicon 2) |
| JT08 | R | AAATCTGGAGCCGGTGAG | 18 | pJT_pTF-mCherry_MS hit-EYFP  (backbone amplicon 2) |
| JT05 | F | CGCTCACCGGCTCCAGATTTATCAGCAATAAACCAGCC | 38 | pJT_pTF-mCherry_MS hit-EYFP  (backbone amplicon 3) |
| JT02 | R | GTCGCTTCACGTTCGCTC | 18 | pJT_pTF-mCherry_MS hit-EYFP  (backbone amplicon 3) |
| JT09 | F | GATACGCGAGCGAACGTGAAGCGACCCTCGAGCTTGAGTCTCTG | 44 | pJT_pTF-mCherry_MS hit-EYFP  (*pTF-mCherry* cassette) |
| JT18 | R | CGAAACACGGAAACCGAAGACC | 22 | pJT_pTF-mCherry_MS hit-EYFP  (*pTF-mCherry* cassette) |

**Table S3 (continued):** Molecular cloning primers.

| **Primer ID** | **Forward [F]**  **or Reverse [R]** | **Sequence**  **[5’ → 3’]** | **Length**  **[nt]** | **Corresponding vector**  **and amplicon details** |
| --- | --- | --- | --- | --- |
| JT19 | F | TTCGGTTTCCGTGTTTCGGAACCCCACCCTTCGTCC | 36 | pJT_pTF-mCherry_MS hit-EYFP  (*Phatr3_J23658* promoter) |
| JT20 | R | TCACGTTCTCGATTATCG | 18 | pJT_pTF-mCherry_MS hit-EYFP  (*Phatr3_J23658* promoter) |
| JT21 | F | ATGGTGAGCAAGGGCGAGG | 19 | pJT_pTF-mCherry_MS hit-EYFP  (*EYFP*) |
| JT22 | R | TTTAGAGTTTTACTTGTACAGCTCGTCC | 28 | pJT_pTF-mCherry_MS hit-EYFP  (*EYFP*) |
| JT23 | F | TACAAGTAAAACTCTAAAGGAGCTTC | 26 | pJT_pTF-mCherry_MS hit-EYFP  (*Phatr3_J23658* terminator) |
| JT24 | R | CCGCGTTTCCAGACTTTACTACTTGTCATACCGAATTTAG | 40 | pJT_pTF-mCherry_MS hit-EYFP  (*Phatr3_J23658* terminator) |
| JT25 | F | CGATAATCGAGAACGTGAATGTTTCGTTTCTCTGCCGTG | 39 | pJT_pTF-mCherry_pTF.CREG1-EYFP  (*pTF.CREG1*) |
| JT26 | R | CTCGCCCTTGCTCACCATAAGACGTTCCAACTGCATATC | 39 | pJT_pTF-mCherry_pTF.CREG1-EYFP  (*pTF.CREG1*) |
| JT27 | F | CGATAATCGAGAACGTGAATGATTCCTCCAGCAACAAGC | 39 | pJT_pTF-mCherry_pTF.CatCh1-EYFP  (*pTF.CatCh1*) |
| JT28 | R | CTCGCCCTTGCTCACCATCACCATTGGTATGTTCGACGC | 39 | pJT_pTF-mCherry_pTF.CatCh1-EYFP  (*pTF.CatCh1*) |
| JT29 | F | CGATAATCGAGAACGTGAATGAAACAGCACCTCTCCC | 37 | pJT_pTF-mCherry_pTF.ap1-EYFP  (*pTF.ap1*) |
| JT30 | R | CTCGCCCTTGCTCACCATAATCGTATCTTGTAAGCGAAGC | 40 | pJT_pTF-mCherry_pTF.ap1-EYFP  (*pTF.ap1*) |
| JT31 | F | GCTAACAGGAGGAATTAACCATGTACCCCTCCCTGCGCCAT | 41 | PtpBAD-CTHF expression vector (*Δ18_pTF.CREG1*) |
| JT32 | R | CTACCGCGAGGGACCAGCGCAAGACGTTCCAACTGCATATCAGGAGAAAGCTTAG | 55 | PtpBAD-CTHF expression vector  (all truncations) |
| JT33 | F | GCTAACAGGAGGAATTAACCATGGAGCTTGATAAACAATCTGATGAATGCCCG | 53 | PtpBAD-CTHF expression vector  (*Δ26_pTF.CREG1*) |
| JT34 | F | GCTAACAGGAGGAATTAACCATGTCTGATGAATGCCCGTGTGAACCTCTACAA | 53 | PtpBAD-CTHF expression vector  (*Δ31_pTF.CREG1*) |
| JT35 | F | GCTAACAGGAGGAATTAACCATGCTACAACGCCCTGATCGTTTCAACAAAGAGG | 54 | PtpBAD-CTHF expression vector  (*Δ39_pTF.CREG1*) |
| JT36 | F | GCTAACAGGAGGAATTAACCATGCGTTTCAACAAAGAGGAGCTAGCTCGCT | 51 | PtpBAD-CTHF expression vector  (*Δ43_pTF.CREG1*) |

**Table S3 (continued):** Molecular cloning primers.

| **Primer ID** | **Forward [F]**  **or Reverse [R]** | **Sequence**  **[5’ → 3’]** | **Length**  **[nt]** | **Corresponding vector**  **and amplicon details** |
| --- | --- | --- | --- | --- |
| JT37 | F | AAACCAATTGTCCATATTGCATCAGACATTGCC | 33 | PtpBAD-CTHF expression vector (colony PCR primer) |
| JT38 | R | CTTTCGTTTTATTTGATGCCTGGCAGTTCC | 30 | PtpBAD-CTHF expression vector (colony PCR primer) |

**Table S4:** Constructed *Phaeodactylum tricornutum* episomes and *Escherichia coli* vectors with expression cassette details and fusion protein amino acid sequences.

| **Vector ID** | **ORF description** | **5’ UTR** | **Promoter** | **Terminator** | **3’ UTR** | **Protein size [kDa]** | **Role in the**  **study** |
| --- | --- | --- | --- | --- | --- | --- | --- |
| pJT_NR_pTF-AP2 | *pTF* (3 exons, 2 introns), *link* (KGSGSTSGSG), *APEX2*  (*T. pseudonana* codons) | *nitrate reductase* (*NR*) | *NR* | *NR* | *NR* | 84.73 | • pTF localization  • proximity proteomics |
| pJT_native_pTF-mCherry | *pTF* (3 exons, 2 introns), *mCherry* | / | *pTF* | *pTF* | / | 83.75 | • pTF localization  • MDY-64 labeling |
| pJT_pTF-mCherry_pTF.CREG1-EYFP | ORF1: *pTF* (3 exons, 2 introns), *mCherry*  ORF2: *pTF.CREG1* (no introns), *EYFP* | / \| / | *pTF* \| *flavodoxin* (*Phatr3_J23658*) | *pTF* \| *flavodoxin* (*Phatr3_J23658*) | / \| / | 83.75 \| 53.32 | pTF and pTF.CREG1 co-expression |
| pJT_pTF-mCherry_pTF.CatCh1-EYFP | ORF1: *pTF* (3 exons, 2 introns), *mCherry*  ORF2: *pTF.CatCh1* (1 intron*), *EYFP* | / \| / | *pTF* \| *flavodoxin* (*Phatr3_J23658*) | *pTF* \| *flavodoxin* (*Phatr3_J23658*) | / \| / | 83.75 \| 64.41 | pTF and pTF.CatCh1 co-expression |
| pJT_pTF-mCherry_pTF.ap1-EYFP | ORF1: *pTF* (3 exons, 2 introns), *mCherry*  ORF2: *pTF.ap1* (no introns), *EYFP* | / \| / | *pTF* \| *flavodoxin* (*Phatr3_J23658*) | *pTF* \| *flavodoxin* (*Phatr3_J23658*) | / \| / | 83.75 \| 69.33 | pTF and pTF.ap1  co-expression |
| pJT_Δ31_pTF.CREG1-His6** | *pTF.CREG1* (with 31 aa N-terminal deletion), *His6* | / | *pBAD* (arabinose-inducible) | *rrnB* | / | 25.26 | pTF.CREG1 *in vitro* enzymatic assays |

*Likely due to gDNA contamination during RNA extraction and cDNA synthesis (a different clone had a gene version with a spliced intron).

**Only the vector used for large scale purification that led to enzymatic assays is described here in full. All other vectors differed from this one only in the exact pTF.CREG1 truncation and/or the His6 position.

Fusion protein amino acid sequences

>**pTF**-*link*-APEX2

**MFSFKSFVFAALLSSCEASVRARKLSNELIAGFEPRTVVTDHNAIDLDQAKIEDLLATPSTDTFSSARDVYEQGSHSKSFAVLTLDPALTVAVPEGTVISGKNEDGSTVSGTALKSYSVGDTTIEIQYSTGNTQATYVDCQVGGNPDPNTSGCFAANGTVTISGSSGSLPYSYDPLVNNDNGRTIQGFSTAAETRFRPSGGGELFPDFQKFFDYYGTLTYADEWVQSAFARRSTSFSKGDADFSKYGDDGILQAVKKGTAYISIAMYVIRELEDALDDCDRGCDTADCNDDAVHALDEAVAFWTGSLEGKDGSGSGVLMYGLADTRCINFKTCGRNGDEASGKSKVNFEIFRHFDTMQRELTNKRCNSARATKEKIVPWMIVPLIQGVLRYAYIIENDGFTEKAEAEGATFAAAVLPYVASCNAEDAKTIYDNMRVGLGGGASYSTVFDAFVRQSSCMGISCSDIGGYWNEAQGAYEEGAGPCGGSGSGDDGANVGLAVGLSIGGLLVLVLLGVVLRRRRSASGSTVEFKDTGSTPI***KGSGSTSGSG*GKSYPTVSADYQDAVEKAKKKLRGFIAEKRCAPLMLRLAFHSAGTFDKGTKTGGPFGTIKHPAELAHSANNGLDIAVRLLEPLKAEFPILSYADFYQLAGVVAVEVTGGPKVPFHPGREDKPEPPPEGRLPDPTKGSDHLRDVFGKAMGLTDQDIVALSGGHTIGAAHKERSGFEGPWTSNPLIFDNSYFTELLSGEKEGLLQLPSDKALLSDPVFRPLVDKYAADEDAFFADYAEAHQKLSELGFADA

>**pTF**-mCherry

**MFSFKSFVFAALLSSCEASVRARKLSNELIAGFEPRTVVTDHNAIDLDQAKIEDLLATPSTDTFSSARDVYEQGSHSKSFAVLTLDPALTVAVPEGTVISGKNEDGSTVSGTALKSYSVGDTTIEIQYSTGNTQATYVDCQVGGNPDPNTSGCFAANGTVTISGSSGSLPYSYDPLVNNDNGRTIQGFSTAAETRFRPSGGGELFPDFQKFFDYYGTLTYADEWVQSAFARRSTSFSKGDADFSKYGDDGILQAVKKGTAYISIAMYVIRELEDALDDCDRGCDTADCNDDAVHALDEAVAFWTGSLEGKDGSGSGVLMYGLADTRCINFKTCGRNGDEASGKSKVNFEIFRHFDTMQRELTNKRCNSARATKEKIVPWMIVPLIQGVLRYAYIIENDGFTEKAEAEGATFAAAVLPYVASCNAEDAKTIYDNMRVGLGGGASYSTVFDAFVRQSSCMGISCSDIGGYWNEAQGAYEEGAGPCGGSGSGDDGANVGLAVGLSIGGLLVLVLLGVVLRRRRSASGSTVEFKDTGSTPI**MVSKGEEDNMAIIKEFMRFKVHMEGSVNGHEFEIEGEGEGRPYEGTQTAKLKVTKGGPLPFAWDILSPQFMYGSKAYVKHPADIPDYLKLSFPEGFKWERVMNFEDGGVVTVTQDSSLQDGEFIYKVKLRGTNFPSDGPVMQKKTMGWEASSERMYPEDGALKGEIKQRLKLKDGGHYDAEVKTTYKAKKPVQLPGAYNVNIKLDITSHNEDYTIVEQYERAEGRHSTGGMDELYK

>**pTF.CREG1**-EYFP

**MFRFSAVILYAMLTLSGAYPSLRHPFELDKQSDECPCEPLQRPDRFNKEELARWMVHSMDWGVLTTISTRLPDGQPFGNVYSFVDGPCGKGMGTPYFYGTYMDQSFHDSKQNDKVSFTLTEASLPSVCHAGASKDCSISHANAGDPESPVCARLTLSGKLVEVNPDSEEYTRAQAAFFQRHPQMATWPSEHNWIIAKLEIEDLWLINFYGGAAILSIEEYFSAKLSPDMQLERL**MVSKGEELFTGVVPILVELDGDVNGHKFSVSGEGEGDATYGKLTLKFICTTGKLPVPWPTLVTTFGYGLQCFARYPDHMKQHDFFKSAMPEGYVQERTIFFKDDGNYKTRAEVKFEGDTLVNRIELKGIDFKEDGNILGHKLEYNYNSHNVYIMADKQKNGIKVNFKIRHNIEDGSVQLADHYQQNTPIGDGPVLLPDNHYLSYQSALSKDPNEKRDHMVLLEFVTAAGITLGMDELYK

>**pTF.CatCh1**-EYFP

**MIPPATSMAFFAFVLLPSYGAYYASAEVVKFGACMNEPTGQYRCARNSTYCDDAELWQAPWQLDDVQGRIVCGCQDAHVGACFPSHGHLSVCRAEAAACPSGEGFSDLARFYNDGKTCMCHGLGEHPYSDIVLEFEETKYGACVIDASTTRCVMHSYMCEAGQSWLPPDILSTRRSEECFCHEVRTGACSHPLGSICAVDADSCQEDSTFLSPAETVAQGLDCRLCAPADVQLTQAGGVDQDAPAPTPTLGNPPTPPSNPYEDFSPHSAPQDTITGSTNENLSSSSSSGMGIRKPAFVALTVLAVLSVLANVVLFRKLRRRNQDKREATGPVTKEEVEKDSAASNIPMV**MVSKGEELFTGVVPILVELDGDVNGHKFSVSGEGEGDATYGKLTLKFICTTGKLPVPWPTLVTTFGYGLQCFARYPDHMKQHDFFKSAMPEGYVQERTIFFKDDGNYKTRAEVKFEGDTLVNRIELKGIDFKEDGNILGHKLEYNYNSHNVYIMADKQKNGIKVNFKIRHNIEDGSVQLADHYQQNTPIGDGPVLLPDNHYLSYQSALSKDPNEKRDHMVLLEFVTAAGITLGMDELYK

>**pTF.ap1**-EYFP

**MKQHLSLLILFSCLRAAFSVERLCSNPDTGVFIGLAVDEKVYVSVPGGPDPDTGKIQSWNLVDEDSIGPSHSFTGRYLQVVPDEGRVYPSRGEHLTDLKQLENNSPYVAFRLDVSKEGAGWHTLFLRWTGGDTVGGGDSVFVSLHSLSKSKSAIAGHRSLKPLKVPIDSTFRNFAGCCYDMQTHACPCQKSQPTNETCPNYIVREEAAKFGAQCSVGPGVMEFVDSPQWYLYAGQEVGNVMDFDAEPWDATCEAEGSDTADSGRDYASWNIPKGEFELRIYAREDGTAVDAIYIAGPNGKAPGLLQKFGVGDSTICTRSGSSGLGSALRTSGFILLIAGVVLGALVAVAKTDQGSAAMHEVLQHISQRRVPNPSVSSDGIMNTYSELRLQDTI**MVSKGEELFTGVVPILVELDGDVNGHKFSVSGEGEGDATYGKLTLKFICTTGKLPVPWPTLVTTFGYGLQCFARYPDHMKQHDFFKSAMPEGYVQERTIFFKDDGNYKTRAEVKFEGDTLVNRIELKGIDFKEDGNILGHKLEYNYNSHNVYIMADKQKNGIKVNFKIRHNIEDGSVQLADHYQQNTPIGDGPVLLPDNHYLSYQSALSKDPNEKRDHMVLLEFVTAAGITLGMDELYK

>**Δ31_pTF.CREG1**-His6

**SDECPCEPLQRPDRFNKEELARWMVHSMDWGVLTTISTRLPDGQPFGNVYSFVDGPCGKGMGTPYFYGTYMDQSFHDSKQNDKVSFTLTEASLPSVCHAGASKDCSISHANAGDPESPVCARLTLSGKLVEVNPDSEEYTRAQAAFFQRHPQMATWPSEHNWIIAKLEIEDLWLINFYGGAAILSIEEYFSAKLSPDMQLERL**ALVPRGSHHHHHHDYKDDDDK

**PREDICTED SIGNAL PEPTIDE**
