## Supplementary material for "Proximity proteomics in a marine diatom reveals a putative cell surface-to-chloroplast iron trafficking pathway": Key Resources Table

| **Key Resources Table** | | | | |
| --- | --- | --- | --- | --- |
| **Reagent type (species) or resource** | **Designation** | **Source or reference** | **Identifiers** | **Additional information** |
| gene  (*Phaeodactylum tricornutum*) | *pTF*; *ISIP2a* | PMID: 29539640 | Ensembl-Protists:Phatr3_J54465 | Jeffrey B. McQuaid |
| gene  (*Phaeodactylum tricornutum*) | *pTF.CREG1* | This paper | Ensembl-Protists:Phatr3_J51183 | The first subcellular localization and enzymatic activity reports for this *P. tricornutum* protein |
| gene  (*Phaeodactylum tricornutum*) | *pTF.CatCh1* | This paper | Ensembl-Protists:Phatr3_J52498 | The first subcellular localization report for this *P. tricornutum* protein |
| gene  (*Phaeodactylum tricornutum*) | *pTF.ap1* | This paper | Ensembl-Protists:Phatr3_J54986 | The first subcellular localization report for this *P. tricornutum* protein |
| gene  (*Glycine max*) | *APEX2* | PMID: 25419960 | NCBI-Gene:553156 | APEX2 mutations relative to wild-type APX (encoded by NCBI-Gene:553156): K14D, W41F, E112K, A134P |
| strain, strain background (*Phaeodactylum tricornutum*) | *Phaeodactylum tricornutum*; *P. tricornutum*; WT *P. tricornutum* | NCMA | Catalog #:CCMP632 Starter Culture 2x15ml | CCMP632 is synonymous with CCMP2561 and CCAP 1055/1 |
| strain, strain background (*Escherichia coli*) | EPI300 | Lucigen | Catalog #:EC300110 | Electrocompetent; recommended for bacterial conjugation of diatoms |
| strain, strain background (*Escherichia coli*) | BL21 | NEB | Catalog #:C2530H | Chemically competent |
| genetic reagent (*Phaeodactylum tricornutum*) | *ΔpTF P. tricornutum* | PMID: 29539640 |  | Jeffrey B. McQuaid; TALEN-generated pTF knockout *P. tricornutum* strain;  available from the Allen Lab |
| transfected construct (*Phaeodactylum tricornutum*) | pJT_NR_pTF-AP2 | This paper |  | See ***Figure 2B***, **Materials and Methods**, and **Supplemental file 1—Table S4**; available from the Allen Lab |
| transfected construct (*ΔpTF Phaeodactylum tricornutum*) | pJT_native_pTF-mCherry | This paper |  | See **Materials and Methods** and **Supplemental file 1—Table S4**;  available from the Allen Lab |
| transfected construct (*Phaeodactylum tricornutum*) | pJT_pTF-mCherry  _pTF.CREG1-EYFP | This paper |  | See **Materials and Methods** and **Supplemental file 1—Table S4**;  available from the Allen Lab |
| transfected construct (*Phaeodactylum tricornutum*) | pJT_pTF-mCherry  _pTF.CatCh1-EYFP | This paper |  | See **Materials and Methods** and **Supplemental file 1—Table S4**;  available from the Allen Lab |
| transfected construct (*Phaeodactylum tricornutum*) | pJT_pTF-mCherry  _pTF.ap1-EYFP | This paper |  | See **Materials and Methods** and **Supplemental file 1—Table S4**;  available from the Allen Lab |
| transfected construct (*Escherichia coli* BL21) | pJT_Δ31_pTF.CREG1-His6 | This paper |  | See **Materials and Methods** and **Supplemental file 1—Table S4**;  available from the Allen Lab |
| antibody | anti-pTF (rabbit monoclonal) | PMID: 29539640 |  | WB (1:10,000 of 1.14 mg/mL stock); custom-made at OriGene;  available from the Allen Lab |
| recombinant DNA reagent | pPtPBR1 | Addgene | Catalog #:80388 | Episomal vector for bacterial conjugation of *P. tricornutum*;  available from the Allen Lab |
| recombinant DNA reagent | pTA-Mob | PMID: 24595202 |  | Mobilization plasmid for bacterial conjugation of diatoms;  available from the Allen Lab |
| recombinant DNA reagent | PtpBAD-CTHF | PMID: 30262498 |  | *Escherichia coli* protein expression vector;  available from the Allen Lab |
| sequence-based reagent | JT01 | This paper | PCR primer | CGAATCAGGATCTAAAATGAACGCACGTCTGCGACCTGAGCAA;  see **Materials and Methods** and **Supplemental file 1—Table S3** |
| sequence-based reagent | JT02 | This paper | PCR primer | GTCGCTTCACGTTCGCTC;  see **Materials and Methods** and **Supplemental file 1—Table S3** |
| sequence-based reagent | JT03 | This paper | PCR primer | GATACGCGAGCGAACGTGAAGCGACTCACGTAGTGAAGTGATGTTG;  see **Materials and Methods** and **Supplemental file 1—Table S3** |
| sequence-based reagent | JT04 | This paper | PCR primer | TTCCAGACGTAGAACCACTCCCTTTGATAGGAGTGCTGCCAGTG;  see **Materials and Methods** and **Supplemental file 1—Table S3** |
| peptide, recombinant protein | PrimeSTAR GXL DNA Polymerase | Takara Bio | Catalog #:R050B | Recommended for amplifying (parts of) diatom episomal vectors |
| peptide, recombinant protein | Bovine Serum Albumin, Biotinylated | Thermo Fisher Scientific | Catalog #:29130 |  |
| peptide, recombinant protein | Streptavidin, horseradish peroxidase (HRP) conjugate | Thermo Fisher Scientific | Catalog #:S911 | WB (1:15,000) |
| peptide, recombinant protein | WesternSure® Pre-stained Chemiluminescent Protein Ladder | Li-COR | Catalog #:926-98000 | Recommended for streptavidin blotting |
| commercial assay or kit | Phire Plant Direct PCR Master Mix | Thermo Fisher Scientific | Catalog #:F160L | Recommended for diatom genotyping |
| commercial assay or kit | Phire Plant Direct PCR Kit | Thermo Fisher Scientific | Catalog #:F130WH | Recommended for diatom genotyping |
| commercial assay or kit | WesternBreeze™ Chemiluminescent Kit, anti-rabbit | Thermo Fisher Scientific | Catalog #:WB7106 |  |
| commercial assay or kit | TMT10plex™ Isobaric Label Reagent Set | Thermo Fisher Scientific | Catalog #:90406 |  |
| chemical compound, drug | Amplex™ UltraRed Reagent | Thermo Fisher Scientific | Catalog #:A36006 |  |
| chemical compound, drug | 3,3’-Diaminobenzidine (DAB) | Sigma-Aldrich | Catalog #:D8001 |  |
| chemical compound, drug | D-(+)-Biotin-tyramine amide (biotin-phenol) | Berry & Associates | Catalog #:BT 1015 |  |
| software, algorithm | SEQUEST (v. 28, rev. 12) algorithm | PMID: 24226387 |  | Algorithm for matching tandem mass spectra with peptide sequences |
| software, algorithm | JMP | SAS Institute |  | Interactive statistical discovery software |
| software, algorithm | R | R Core Team |  | Free software environment for statistical computing and graphics |
| software, algorithm | ASAFind | PMID: 25438865 |  | Plastidial protein localization prediction tool for algae with red secondary plastids; available at <https://rocaplab.ocean.washington.edu/tools/asafind/> |
| software, algorithm | BLASTP | PMID: 2231712 |  | Algorithm for identifying homologous proteins |
| software, algorithm | HMMER | PMID: 9918945 |  | Software for identifying homologous proteins (or nucleotide sequences) using profile hidden Markov models |
| software, algorithm | MAFFT | PMID:  12136088 |  | Multiple amino acid sequence alignment algorithm based on fast Fourier transform |
| software, algorithm | SeaView 4 | PMID:  19854763 |  | Graphical User Interface (GUI)-based molecular phylogeny software |
| software, algorithm | IQ-TREE | PMID:  25371430 |  | Algorithm for inferring phylogenetic trees by maximum likelihood |
| software, algorithm | Phyre2 | PMID: 25950237 |  | Web server to predict and analyze protein structure, function and mutations; available at <http://www.sbg.bio.ic.ac.uk/~phyre2/html/page.cgi?id=index> |
| software, algorithm | UCSF Chimera version 1.11.1 | PMID: 15264254 |  | Software for interactive visualization and analysis of molecular structures and related data |
| other | Yeast Vacuole Membrane Marker MDY-64 | Thermo Fisher Scientific | Catalog #:Y7536 | Membrane stain |
| other | Pierce Streptavidin Magnetic Beads | Thermo Fisher Scientific | Catalog #:88816 | Protein-coated iron oxide microparticles |
| other | *Phaeodactylum tricornutum* proteome | UniProt | Proteome-ID:UP000000759 | Reference proteomic database |
| other | Phatr3 *P. tricornutum* genomic database | Ensembl Protists | Genome-Assembly:ASM15095v2 | Reference genomic database |
| other | Transcriptomic data | PMID: 27973599 | S1 Dataset. Active transcriptome and assignment of genes to WGCNA modules and response types. | *P. tricornutum* transcriptomic dataset; see ***Figure 5—figure supplement 1—source data 1*** |
| other | Marine Microbial Eukaryote Transcriptome Sequencing Project (MMETSP) database | PMID: 24959919 | NCBI-BioProject:PRJNA231566 | Transcriptomic database |
